# Quantifying bristle cell organization in *Drosophila melanogaster* using spatial clustering features

**DOI:** 10.1101/2025.07.29.667315

**Authors:** Wisdom K. Attipoe, Audrey Neighmond, Oakley Waters, S.H Dinuka Sewwandi De Silva, Ginger L. Hunter, Emmanuel Asante-Asamani

**Affiliations:** Department of Mathematics, Clarkson University, Potsdam, NY 13699; Department of Biology, Howard University, Washington, DC 20059; Department of Biology, Clarkson University, Potsdam, NY, 13699; Department of Mathematics, University of Kelaniya, Kelaniya, Sri Lanka

**Keywords:** Tissue patterning, *Drosophila melanogaster*, DBSCAN, Machine Learning, Clustering

## Abstract

Developing tissues reproduce nearly identical patterns from animal to animal, despite the process being stochastic at the cell level. A major challenge for researchers is the ability to quantify and classify complex cell and tissue spot patterns across wild type and perturbed conditions. Here, we use the organization of small sensory bristles on the fruit fly thorax as a model system to address this problem. A well-known and easily observable distinguishing feature of spot patterns is density. Beyond this, it is unclear how to quantitatively distinguish between patterns in a reproducible way. Our work evaluates the utility of the spatial clustering of spot patterns in quantifying their organization. We propose four clustering features, obtained using the density-based spatial clustering of applications with noise (DBSCAN) algorithm. Together with pattern density we assess how these features can quantify and distinguish between bristle patterns in a variety of wild-type and mutant flies to confirm known perturbations and discover new ones. Our results show that a combination of spot pattern density and the variance between the size of clusters is sufficient to distinguish between 70% of patterns from wild-type and mutant tissues.

## 1 Introduction

Repeating spot patterns are found throughout our world, both man-made and in nature. It is easy to look at a spot pattern and have a sense of its organization - regular, irregular, more or less dense-but what is more difficult is to reproducibly quantify how any two patterns are similar or different. The array of small sensory bristles on the dorsal thorax of the fruit fly Drosophila melanogaster is a model system for the study of biological spot patterns. These bristles enable the fly to sense its environment as part of the peripheral nervous system. The tissue-wide organization of this bristle array is similar among wild-type Drosophila melanogaster, yet each individual is distinct. This is because the signaling mechanisms that govern the selection of bristle cells from an initially bipotential epithelium are stochastic. Therefore, at the onset of patterning it is not possible to tell which cells will acquire the sensory bristle fate and which will not.

Patterning of the adult Drosophila notum sensory bristle array is primarily driven by Notch-mediated lateral inhibition [1]. All cells in the initially unpatterned epithelium choose either a bristle precursor cell or epithelial cell fate according to their Notch activation levels: high levels of activated Notch are associated with epithelial fate, while low levels are associated with bristle precursor fate. The mechanisms of Notch signaling have been reviewed in depth [2, 3]. Briefly, Delta activates Notch in trans, leading to cleavage of the Notch intracellular domain and its translocation to the nucleus, where it can participate in regulating gene expression, including repression of *delta*. In order to achieve the wild-type organization of bristles, lateral inhibition takes place between both adjacent neighboring cells as well as more distant neighbors (i.e., +1 cell diameter away)[4, 5, 6]. Long-range signaling is achieved in part by the formation of actin-rich cytonemes on the basal surface of all cells [5, 7]. Disrupted cytoneme-mediated signaling leads to the formation of bristle patterns that are more dense [5, 8, 9, 10]. Studies of the patterning notum have led to numerous insights into the mechanisms of Notch signaling, planar cell polarity, and tissue homeostasis [11, 12]. The developing and adult bristle array has been the subject of many genetic screens and other studies of cell signaling [13, 14]. Many of these studies score for strong phenotypes, for example, transformation of bristle support cell fates, loss of bristles, gain of bristles, or failure to form the notum. More subtle changes to bristle organization are more difficult to capture. One reason for this arises from the distinctiveness of each animal’s bristle pattern: what is the range of wild-type patterning? How can we detect if a perturbation to the overall pattern lies outside the wild-type range? Another reason is due to the lack of tools to quantify complex changes in tissue-wide spot patterns. Local changes are easier to determine, and quantified by several means, including the distance between pairs of bristles cells [5]. However, local measurements cannot capture the overall organizational differences across tissues.

One possible tissue-wide organization is the formation of groups of bristle cells due to differences in the spacing between pairs of bristles cells at different locations in the tissue. This could result from location specific differences in the strength of lateral inhibition and Notch-Delta signaling. Cluster analysis is a very common tool in machine learning for identifying and characterizing inherent groups of related points within a point cloud [15]. The most common clustering algorithm is K-means, which organizes points based on their proximity to each other into predetermined groups [16]. Popular distance metrics such as Euclidean distance or Manhattan distance tend to place clusters within a rectangle or disk, limiting the detection of clusters with more complex geometries, such as bows or rings of different radii. K-means also requires apriori knowledge of the number of clusters in the point cloud thus biasing the search process and forces every point to belong to a cluster [17].

An alternative clustering framework, which overcomes the limitations of K-means, is Density-Based Spatial Clustering of Applications with Noise (DBSCAN)[18, 19]. DBSCAN uses the local density of points to track and group points, that can be reached from each other by traveling across disks of fixed radii, into a cluster. Points that do not fall into any cluster are described as outliers. The algorithm has been successfully used to quantify the spatial distribution of cells within an epithelium [20], analyze gene expression patterns [21] and analyze the spatial distribution of proteins involved in DNA repair [22]. In this work, we apply DBSCAN to identify clusters within bristle cell patterns and use properties of these clusters to obtain global features for quantifying spot patterns. We compare the distribution of these features in wild-type flies and select mutants suspected to alter the organization of bristle cells and train a support vector machine, using the most significant features, to identify an optimal region for wild-type patterns.

## 2 Materials and Methods

### 2.1 Visualizing bristle cell array

To visualize the developing array of bristle precursor cells, we co-expressed two GFP labels: (1) shotgun-GFP, an endogenously GFP-tagged, full-length E-cadherin that labels the cell boundaries of all cells in the notum; (2) neur-GMCA, which expresses a GFP-tagged, constitutively active fragment of Moesin to label the filamentous actin cytoskeleton [23] under the control of the *neuralized* promoter, which is expressed in bristle precursor cells [24]. To perturb the pattern of sensory bristles on the dorsal thorax, we used the GAL4/UAS system to knockdown genes of interest via RNAi in the pupal notum region during patterning stages [25]. Specifically, we used pannier-GAL4 [26] to express UAS-RNAi responders throughout the central notum. For controls, we expressed UAS-white RNAi or UAS-Dicer2.

### 2.2 Fly strains and husbandry

Fly strains used in this study include the following. **GAL4 drivers**: shotgun-GFP, neur-GMCA/CyOGFP; pannier-GAL4/TM6BTb (Hunter laboratory). **UAS responders**: UAS white RNAi (BL#33644). UAS-Dicer2 (BL#24650). UAS-Dicer2; UAS-Mindbomb1 (BL#27320, Dcr2 added by this study). UAS-Scabrous RNAi (BL#63585). UAS-Myosin XV RNAi (BL#41691). UAS-myosin VIIa RNAi (BL#41690). UAS-myosin VIIb RNAi (BL#41717). UAS-SCAR RNAi (BL#36121). UAS-Myosin VI RNAi (BL#28064) All RNAi fly stocks were acquired from the Bloomington Drosophila Stock Center (BL), Bloomington, IN. Fly stocks were maintained at 18 degrees Celsius on standard fly food. The crosses were kept at room temperature. White pre-pupae of the F1 generation (0 hours after pupariation, hAP) were collected and screened for balancer phenotypes. After this, they were aged for 24 hours at 25 degrees Celcius.

### 2.3 Microscopy

24 hAP pupae were dissected and mounted on slides as previously described [27]. All imaging was performed on a Leica iDM8 confocal microscope (HC PL FLUOTAR 40x/0.80). Z-stacks of 5-10 images were collected with a z-step size of 2 micrometers. At minimum, one heminota per animal was captured, and for most animals, the full notum, centered on the animal midline, was imaged. To generate images to quantify the area of a bristle precursor cell at 24h AP, we used a digital zoom to image a subset of the tissue.

### 2.4 Image processing

Processing of the notum images was performed in FIJI/Image J. For overall spot patterns, images were first adjusted for brightness and contrast. Adjusted images were then blurred (Gaussian) with *σ* = 2.0 to average intensities after which the thresholding and binarization tools were applied to isolate bristle cells as bright spots with a dark background. The image was then rotated to keep the midline in a vertical orientation. The full notum was then cropped into two heminota, using the animal midline as a reference. Finally, the visible area of the heminota as well as the x-y coordinates of bristle cell centroids within the heminota were extracted for further analysis. The image processing workflow is shown in Fig. 1. For quantifying bristle precursor cell area in control-white animals, cells were identified by their shape: daughter cells in the lineage display asymmetrical apical surfaces compared to surrounding epithelial cells. We then manually traced the cells in FIJI/ImageJ to find the apical area of the bristle precursor cells.

**Figure 1:**
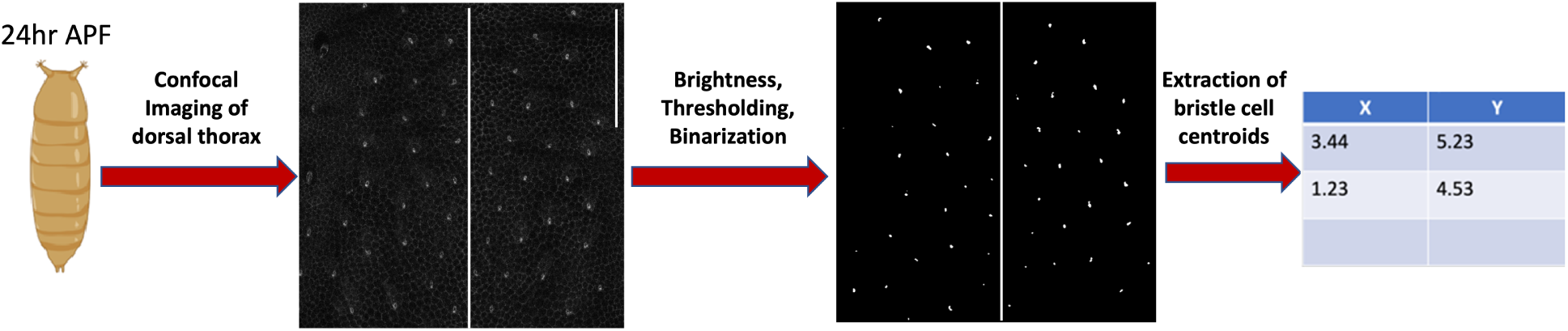
Image processing workflow. Steps for processing microscopy images of fly notum 24 hours after pupae formation (h AP) to obtain density and clustering features. Scale bar on top right hand corner of unprocessed image is 100 microns.

### 2.5 Global features of spot patterns

#### Relative sensory area (RSA)

To quantify the density of the bristle pattern in each notum, we quantified the average area of single bristle cells at 24h AP (24.5 ± 1.29*µm*^2^*, n* = 8*cells, N* = 2*pupae*) and the corresponding number of bristle cells to calculate the total area of the heminota occupied by bristle cells. We refer to this product as the total sensory area. Pattern density is defined as the total sensory area divided by the total area of the heminota (obtained from Image J), which is a measure of the number of bristle cells per unit area of tissue. More precisely, we call this ratio the *Relative Sensory Area* (RSA). Animals with a high uniform density of sensory bristles should have a high RSA value. Note, however, that this feature does not capture the spatial organization of bristle precursor cells, but rather the relative area of the tissue they occupy. Consequently, the RSA can not distinguish between patterns with the same number of bristle cells either packed into a corner of the tissue or spread out uniformly. In order to capture the spatial arrangement of bristle cells, we use spatial clustering to define the following additional features.

#### Clustering features

The clustering features of bristle cell patterns are based on first clustering the centroids of bristle cells and then quantifying the organization of the clusters. The methods are applicable to quantifying the organization of any point cloud, 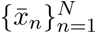 where 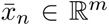 and *m* ∈ Z^+^, with noticeable differences in the local density of points. Thus, we describe the clustering features for a general point cloud 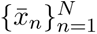. We use the Density-Based Spatial Clustering for Applications with Noise (DBSCAN) but other clustering methods may be used. Our primary motivation for using DBSCAN is the generality in the structure of clusters it can identify.

##### DBSCAN Algorithm

The DBSCAN algorithm initializes *ɛ*-balls around each point and classifiers them as *core points*, *boundary points* or *outliers* according to the following rules. *Core points* will have at least some chosen minimum number of points (MinPts) within their *ɛ*-balls. Points with less than MinPts in their *ɛ*-ball but are contained in the *ɛ*-ball of a core point are called *boundary points*. Any other point is classified as an *outlier*. A group of points form a cluster if any pair of core points can be reached by traveling across intersecting *ɛ*-balls. The major parameters, *ɛ >* 0 and *MinPts* ∈ Z^+^ are chosen to maximize the separation between clusters while minimizing the proximity of points within the same cluster. The Silhouette score for each data point *x̄_n_* is a measure of the likelihood of placing it in the correct cluster. Extending this to the point cloud, we use the mean silhouette score over all data points as a measure of the quality of clustering. We will refer to this as the *pattern silhouette score*, and define it more precisely in later paragraphs. The optimal clustering of bristle cells is thus achieved with values of *ɛ* and *MinPts* that maximize the pattern silhouette score.

Based on the optimal clustering, we quantified the spatial organization of a point cloud using five features. Cluster number (CN), Outlier prevalance (OP), Cluster quality (CQ), Cluster size variance (CSV).

##### Cluster number (CN)

This is the number of clusters identified by the DBSCAN algorithm.

##### Outlier prevalence (OP)

This is the proportion of points in the point cloud that are classified as outliers. Outliers are points which cannot be associated with any cluster, thus far away from other points in the tissue. In our application, OP is indicative of bristle cells with stronger than normal lateral inhibition, thus creating a larger region of epithelial cells around it.

##### Cluster quality (CQ)

is measured using the pattern silhouette score. Mathematically, we define the silhouette score for each data point *x̄_i_* as

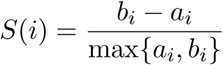

where 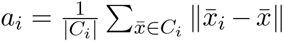 is the average distance between data point *x̄_i_* and other data points *x̄* in its cluster *C_i_*. Here |*C_i_*| denotes the number of points in the cluster *C_i_*. The parameter 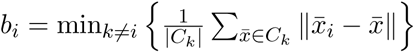 is the average distance between a data point *x̄_i_* and other data points in the nearest cluster. Thus, the Silhouette score for each point *S*(*i*) ∈ [−1, 1], with negative values denoting the degree of misclassifications. The pattern silhouette score (Cluster quality) is then defined as

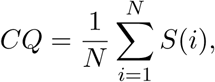

where *N* is the total number of points under consideration.

##### Cluster size variance (CSV)

measures the degree of homogeneity or uniformity in the size of clusters within the pattern. We express it as

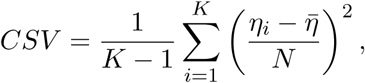

where *η* is the average size of clusters in the pattern, *η_i_* is the number of points in the *i^th^* cluster and *K* is total number of clusters. We make this feature independent of the total number of points by scaling the term (*η_i_* − *η*) by *N* .

### 2.6 Statistical analysis and machine learning

#### Univariate analysis

Kruskal Wallis test, a non-parametric form of ANOVA, was used to determine whether each of the global pattern features described above varied significantly between any fly mutants or controls. The Mann Whitney U test was subsequently used to determine pairwise differences in features between controls and their corresponding mutants. We compared features from Mindbomb1 mutants to Dcr2 and all other mutants to White. All statistical tests were performed at a 5% level of significance.

#### Regression analysis

For mutants that showed a significant difference from their controls in at least one of the features above, we used regression analysis to determine the combination of features most informative in distinguishing each mutant pattern from its control. Patterns were grouped into binary classes with control-0 and mutant-1. Analysis was performed separately for each mutant using logistic regression. For mutants that had a single significant feature, we used the p-value of the regression coefficients to determine the discrimination power of the feature. For mutants with multiple significant features, we used a likelihood ratio test to identify the optimal combination of features. In particular, given the two regression models, *M* 1 : *logit*(*p*) = *β*_0_ +*β*_1_*x*_1_ and *M* 2 : *logit*(*p*) = *β*_0_ +*β*_1_*x*_1_ +*β*_2_*x*_2_ for features *x*_1_*, x*_2_, the likelihood ratio tests investigates the null hypothesis that *β*_2_ = 0. A failure to reject the null hypothesis would suggest that *x*_1_ is sufficient to distinguish the mutant pattern from its control.

#### Support vector machines

We used a subset of features showing the strongest potential to distinguish normal from perturbed patterns (based on our statistical analysis) to train a support vector machine (SVM) for two purposes. First, to identify a range of wild-type patterning and then to quantify the extent to which patterns from each mutation differed from control. In order to identify the range of wild-type patterning, feature information from all the controls (White and Dcr2) were grouped into one class with all mutants, having at least one feature that significantly differed from wild-type, placed into another class. The SVM was trained using 60% of the data and tested on the remaining. To determine the optimal decision boundary, we run a grid search by varying the type of svm kernel (linear, polynomial, radial basis function), as well as the parameters *C* ∈ {0.1, 1, 10, 100, 1000}, *γ* ∈ {0.001, 0.01, 0.1, 1, 10} and *degree* ∈ {2, 3, 4}. In the second application, we constructed a decision boundary that separated data for each mutant from its corresponding control.

## 3 Results

We have developed five global features that can be used to quantify spot patterns. Of particular interest to us are the spot patterns created by bristle cells on the dorsal thorax of fruit flies. Our goal in seeking to quantify these spot patterns is to describe subtle differences in their spatial organization so that wild type patterns can be distinguished from genetic mutations that significantly alter normal bristle cell organization. Our features capture the overall density of the spot pattern as well as the clustering of spots. Our measure of pattern density, referred to as the relative sensory area (RSA), is a measure of the number of cells per unit area of tissue. We developed four clustering features, obtained from analyzing clusters of spots created using the DBSCAN algorithm. They are the cluster number (CN), outlier prevalence (OP), cluster quality (CQ) and cluster size variance (CSV). Cluster number describes the total number of clusters present in the spot pattern. Outlier prevalence defines the fraction of spots that are relatively far away from neighboring spots. Cluster quality and cluster size variance describe the relative separation and compactness of each cluster, and the variance in the size of all clusters respectively. A detailed description of these features has been given in the Material and Methods section.

To evaluate the ability of our proposed global spot pattern features to quantify bristle organization and identify genetic mutations that significantly alter the pattern, we targeted several genes known to perturb the spacing of sensory bristles. Mindbomb1 (Mib1) is an E3 ubiquitin ligase that helps trigger the endocytosis of Delta ligand, a step that is essential for Notch activation [28]. Mib1 mutant flies exhibit an overproduction of sensory bristles [29]. Scabrous is a secreted modulator of Notch signaling whose mutants exhibit an increased density of bristles on the dorsal thorax [30]. The actin cytoskeleton regulator SCAR has been previously shown to contribute to the cytoneme dynamics that support bristle precursor spacing. Loss of SCAR disrupts long-range Notch signaling and thus leads to an increased density of bristles on the dorsal thorax. [5, 9].

We also selected four genes whose effect on bristle patterning are not well understood. The unconventional MyTH4-FERM myosins are actin-based motors involved in the formation and regulation of actin-based cell protrusions, including bristle shafts, filopodia, and cytonemes [31, 32]. The Drosophila genome encodes four MyTH4-FERM myosins: VIIa, VIIb, XV, and XXII. We investigate Myosin VIIa and XV as are both known to play a role in the extension and organization of the sensory bristle shaft [33, 34]. Recently, it was shown that loss of Myosin XV leads to increased final bristle density and transient disorganization during patterning [10]. We also investigate Myosin VIIb and VI, since knockdown of each is associated with bristle specification defects [13]. Although not a MyTH4-FERM myosin, myosin VI is sole minus directed myosin in Drosophila [35]. We did not include Myosin XXII (*myo81F*) in this study. However, for all seven of the genes selected, it is not known how gene knockdown affects other features of bristle organization, beyond density.

We imaged pupae expressing RNAi against these targets at 24 hours after pupariation (hAP) at which time the bulk of the bristle patterning processes are finished. We processed these images to increase visibility of each bristle cell. Details of the image processing is in Materials and Methods and shown in Fig. 1. The notum has a natural symmetry along the animal midline which separates the bristle fields into two lateral halves. Patterns in each half tend to be very similar and would create two obvious clusters. In order to increase the number of patterns collected, identify more intricate clusters, and reduce any noise that the gap around the midline would introduce we split the tissue into two halves (heminota), along the midline, and analyzed the pattern in each heminota separately. This generated 376 patterns across all control and mutant flies. A breakdown of the number of tissues for each fly is given in Table 1.

**Table 1:** Distribution of global pattern features among mutants and controls. Data presented as *Mean* ± *Std*.

| Tissue type | Samples | RSA | CN | OP | CQ | CSV |
| --- | --- | --- | --- | --- | --- | --- |
| White | 70 | $0.021 \pm 0.0059$ | $2.99 \pm 1.27$ | $0.31 \pm 0.23$ | $0.42 \pm 0.11$ | $0.10 \pm 0.10$ |
| Dcr2 | 36 | $0.019 \pm 0.0047$ | $2.89 \pm 1.15$ | $0.38 \pm 0.29$ | $0.41 \pm 0.09$ | $0.09 \pm 0.12$ |
| Mindbomb1 | 44 | $0.039 \pm 0.0097$ | $2.61 \pm 0.84$ | $0.14 \pm 0.09$ | $0.44 \pm 0.10$ | $0.23 \pm 0.17$ |
| Scabrous | 46 | $0.027 \pm 0.0088$ | $3.02 \pm 1.26$ | $0.32 \pm 0.23$ | $0.43 \pm 0.11$ | $0.10 \pm 0.12$ |
| SCAR | 36 | $0.023 \pm 0.0043$ | $2.89 \pm 0.95$ | $0.30 \pm 0.27$ | $0.38 \pm 0.10$ | $0.15 \pm 0.17$ |
| Myosin VIIa | 44 | $0.021 \pm 0.0053$ | $3.05 \pm 1.46$ | $0.33 \pm 0.23$ | $0.38 \pm 0.14$ | $0.12 \pm 0.14$ |
| Myosin XV | 58 | $0.024 \pm 0.0052$ | $2.66 \pm 1.12$ | $0.27 \pm 0.22$ | $0.45 \pm 0.10$ | $0.17 \pm 0.15$ |
| Myosin VIIb | 28 | $0.021 \pm 0.0035$ | $3.25 \pm 1.60$ | $0.35 \pm 0.22$ | $0.37 \pm 0.11$ | $0.13 \pm 0.15$ |
| Myosin VI | 14 | $0.025 \pm 0.0044$ | $2.86 \pm 0.95$ | $0.24 \pm 0.15$ | $0.44 \pm 0.11$ | $0.11 \pm 0.13$ |

For each tissue, we identified the total number of bristle precursor cells, total tissue area, and the *xy*−coordinates of the centroid of each bristle precursor cell. The total tissue area and number of cells were used to calculate the relative sensory area (RSA). The DBSCAN algorithm was used to detect clusters in the *xy*−coordinates for computing our clustering features. In Fig. 2, we show examples of control and mutant patterns along with the clusters we obtained after running DBSCAN. Details on the clustering algorithm and computation of the pattern features can be found in Materials and Methods.

**Figure 2:**
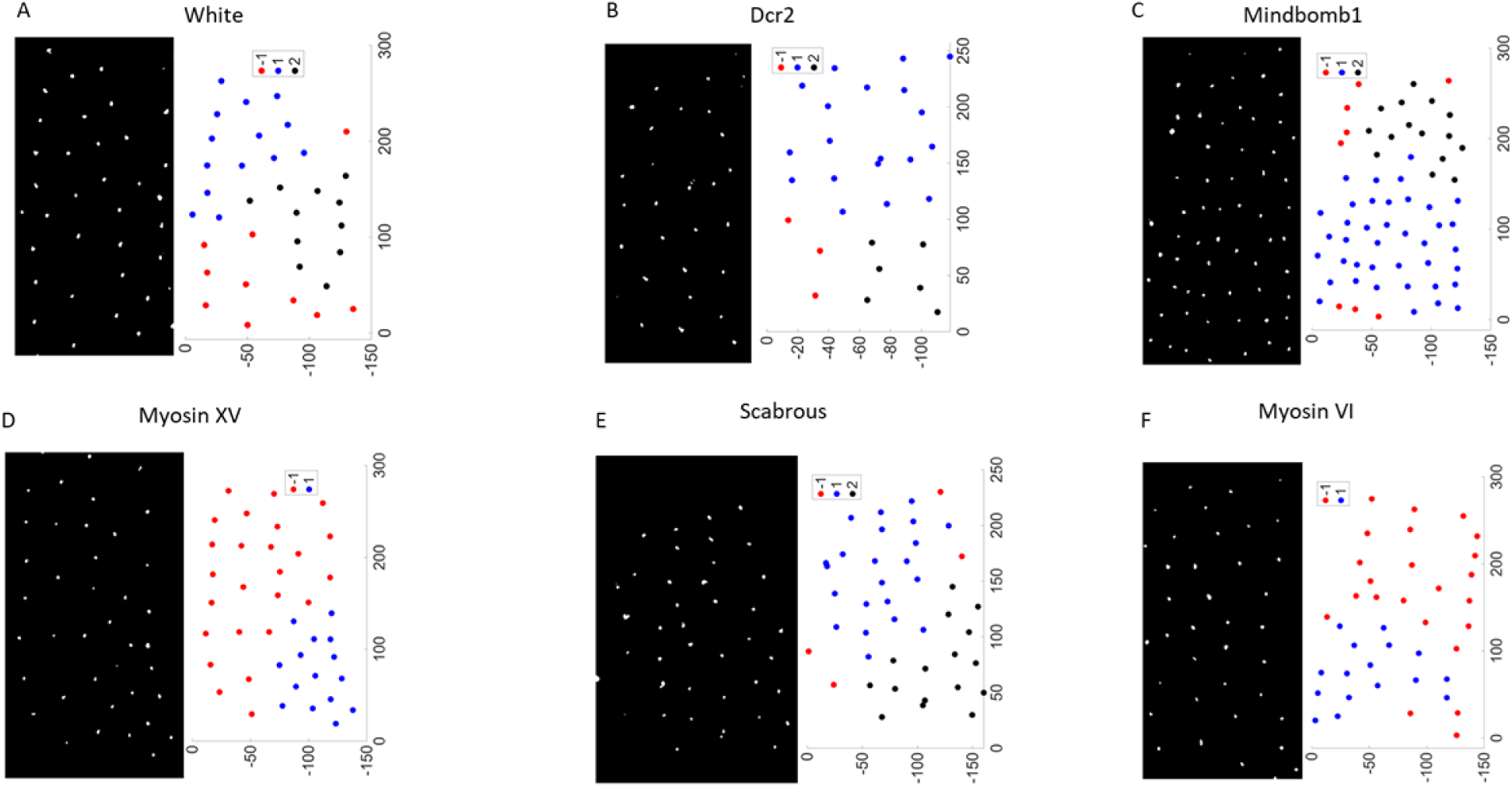
Clustering of bristle precursor cells. Processed images of bristle precursor cells for select mutants and controls with corresponding clustering of centroids using DBSCAN. Colors denote individual clusters of cell centroids. Axes units are in microns.

### 3.1 Ranges for wild type and mutant patterning

Consistent with published literature, we observe that bristle patterns in our control animals (White and Dcr2) are very similar to each other. As can be seen in Table 1 and Fig. 2, bristle cells in control-White tissues formed about 3 clusters with density (RSA) of 0.021 ± 0.006 and a cluster variance (CSV) of 0.10 ± 0.10. The quality of clustering (CQ) measured with the pattern Silhoutte score is 0.42 ± 0.11 with an outlier prevalence (OP) of 0.3, indicating that 30% of bristle precursor cells are distant from major clusters. We did not observe any significant differences between the density or clustering of control-White and control-Dcr2 tissues.

The global features of each mutant were compared to their corresponding controls, using a Mann-Whitney U-test to check for statistically significant differences. Violin plots of the data are shown Fig. 3. Mindbomb1 knockdown patterns were compared with control-Dcr2 while all other mutant patterns were compared to control-White. Our results from Table 1 and Fig. 3 show that Mindbomb1 knockdown patterns have significantly higher density than control (*pvalue <* 0.01), show more variation in the size of clusters (*pvalue <* 0.01) and have a smaller proportion of outliers (*pvalue <* 0.01). However, the number and quality of clusters are not significantly different from control-Dcr2. Myosin XV knockdown patterns also have a higher density than control (*pvalue* = 0.03) and show a larger variation between the size of clusters (*pvalue <* 0.01). But the number and quality of clusters as well as proportion of outliers are not significantly different from wild type. Myosin VI knockdown patterns are much denser than wild type (*pvalue* = 0.014) but similar to wild type in all other features. The only distinguishing feature for Scabrous knockdown tissues is a higher pattern density (*pvalue <* 0.01). SCAR knockdown tissues only show better quality of clustering (*pvalue* = 0.04) without significantly altering any other feature. The remaining knockdown conditions, Myosin VIIa and VIIb, appear to have no significant differences with wild type patterns for the global features under consideration.

**Figure 3:**
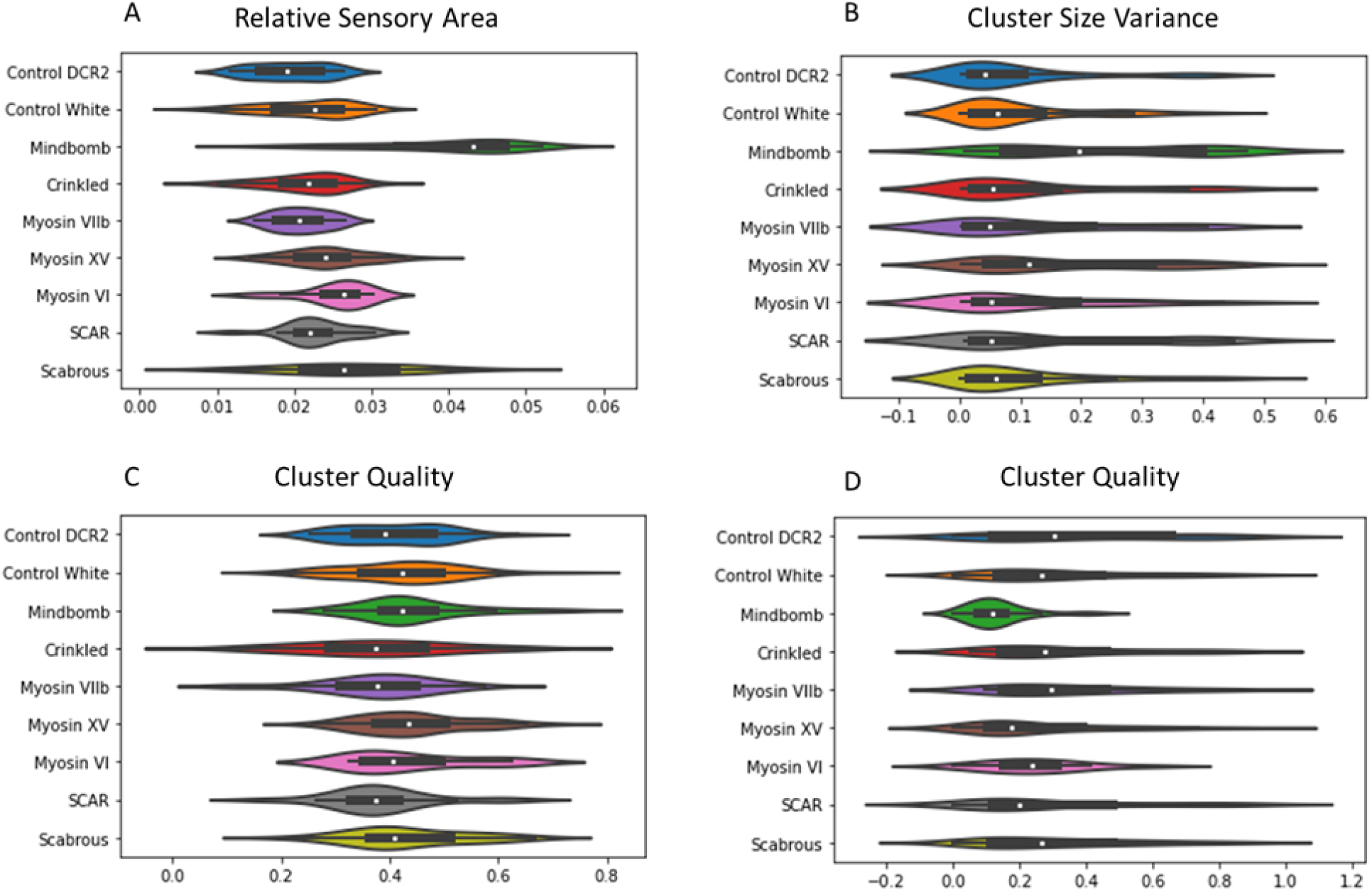
Distribution of global features. Violin plots showing distribution of global pattern features across all heminota.

### 3.2 Optimal combination of features for pattern identification

Next, we wanted to assess the predictive ability of each feature by analyzing its association with the odds of a perturbation in the bristle spot pattern, Also, in mutants where multiple features were identified as significantly different from the respective control, we wanted to identify an optimal combination that will maximize our ability to distinguish perturbed patterns from wild type patterns. Within each mutant population and its corresponding control, we categorized each pattern into a class of either control-0 or mutant-1 and fit logistic regression models in increasing order of complexity (i.e with a single feature, two features, three features etc) and performed a likelihood ratio test. This allowed us to test whether addition of features improves the ability of the model to distinguish between wild type and perturbed patterns. Additionally, we fit a multiple logistic regression model with all features as predictor variables and examined the significance of the regression coefficients. The p-values from the likelihood ratio tests are shown in Table 2 with p-values from multiple logistic regression shown in Table 3.

**Table 2:**
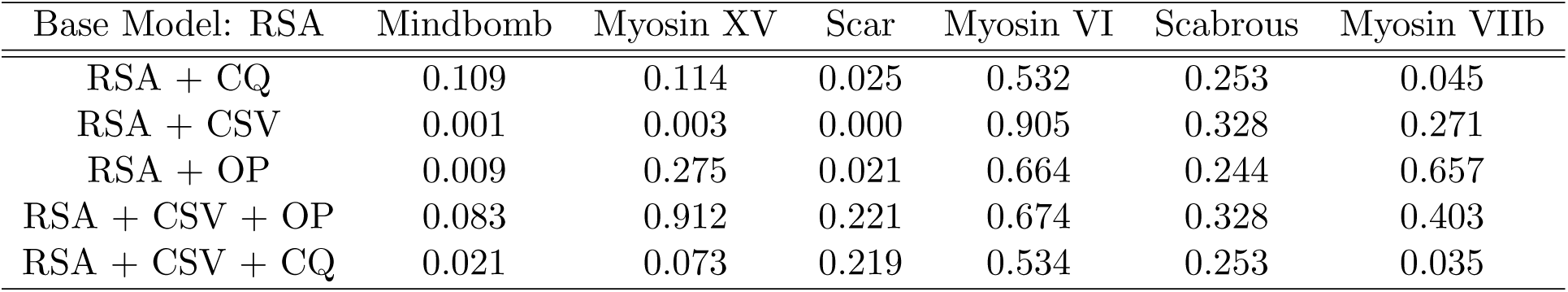
Likelihood ratio test for global features with p-values of corresponding models.

| Base Model: RSA | Mindbomb | Myosin XV | Scar | Myosin VI | Scabrous | Myosin VIIb |
| --- | --- | --- | --- | --- | --- | --- |
| RSA + CQ | 0.109 | 0.114 | 0.025 | 0.532 | 0.253 | 0.045 |
| RSA + CSV | 0.001 | 0.003 | 0.000 | 0.905 | 0.328 | 0.271 |
| RSA + OP | 0.009 | 0.275 | 0.021 | 0.664 | 0.244 | 0.657 |
| RSA + CSV + OP | 0.083 | 0.912 | 0.221 | 0.674 | 0.328 | 0.403 |
| RSA + CSV + CQ | 0.021 | 0.073 | 0.219 | 0.534 | 0.253 | 0.035 |

**Table 3:** Significance of coefficients in multiple logistic regression model indicated by p-values.

| Tissue | RSA | OP | CQ | CSV |
| --- | --- | --- | --- | --- |
| Mindbomb1 | 0.014 | 0.239 | 0.051 | 0.021 |
| Myosin XV | 0.021 | 0.970 | 0.079 | 0.006 |
| Scar | 0.000 | 0.358 | 0.356 | 0.013 |
| Myosin VI | 0.044 | 0.757 | 0.579 | 0.954 |
| Scabrous | 0.000 | 0.254 | 0.199 | 0.475 |
| Myosin VIIb | 0.420 | 0.625 | 0.052 | 0.170 |

Recall from our previous analysis that Mindbomb1 knockdown showed a significant difference in pattern density (RSA), cluster size variance (CSV) and outlier prevalence (OP) when compared with control-Dcr2. Using a base model with just RSA, our likelihood ratio test results (Table 2) indicate that addition of CSV to RSA will likely improve the models predictiveness (*pvalue <* 0.01). Likewise, addition of OP to RSA is likely to improve the models predictiveness (*pvalue* = 0.027). However, results from the multiple logistic regression (Table 3) indicate that RSA and CSV are sufficient to distinguish Mindbomb1 knockdown from control-Dcr2 tissues.

We observed previously that Myosin XV knockdown tissues showed a significant difference in pattern density (RSA) and cluster size variance (CSV) when compared with control-White. A likelihood ratio test indicates that addition of CSV to RSA will likely improve a model’s predictiveness (*pvalue <* 0.01, Table 2). This is supported by a multiple regression model showing both RSA and CSV to have significant regression coefficients (Table 3). We observed a significant difference in cluster quality (CQ) for SCAR knockdown when compared with control-White. However, a single logistic regression model involving CQ as a predictor variable shows only a weak association with the odds of pattern perturbation (*pvalue* = 0.082). Surprisingly, RSA shows a strong association with the odds of perturbation (*pvalue <* 0.01) even though is was not significantly different from control-White (*pvalue* = 0.47). Likewise CSV shows a weak association (pvalue = 0.063) with odds of perturbation although it was not significantly different from control-White (*pvalue* = 0.43). Interestingly, a multiple logistic regression model including all pattern features shows a significant association between both RSA and CSV with the odds of pattern perturbation without a significant association for CQ (pvalue = 0.36, Table 3). Consequently, whereas CQ is significantly different between control-White and SCAR knockdown patterns, it is unlikely to be useful in distinguishing between the two patterns.

Myosin VI knockdown tissues differed significantly from control-White in pattern density. Our likelihood ratio test and regression analysis show that no other feature can improve the predictive ability of RSA in distinguishing wild type from Myosin VI-driven perturbations (Table 2, Table 3). Scabrous knockdown tissues also had significantly higher pattern density compared with control-White. Similar to Myosin VI knockdown, no other feature can improve on the information available from RSA in distinguishing between control-White and Scabrous-driven perturbations (Table 2, Table 3). Whereas no feature was significantly different between control-White and Myosin VIIb, cluster quality appears to have a weak association with the odds of perturbation, as can be seen from the significance of coefficients in the multiple logistic regression as well as likelihood ratio test (Table 2, Table 3). As concluded from our univariate analysis, no feature from Myosin VIIa patterns will be useful in distinguishing it from control-White.

### 3.3 Identification of wild type bristle patterns

The primary focus of our work is to define a range of global features that distinguishes wild type patterns from perturbed ones. Our statistical analysis above has identified three pattern features, density (RSA), cluster size variance (CSV) and cluster quality (CQ), with the potential to classify wild type patterns. In Fig. 4 we show a scatter plot of mutant tissues with a pair of distinguishing features. Note that Mindbomb1 knockdown tissues show a clear separation from control-Dcr2, whereas the separation between the others and control-White is more modest. Note also that Myosin VI and Scabrous knockdown tissues are mostly separated by pattern density (RSA), agreeing with our statistical analysis which found that CSV is not a significant distinguishing feature. Myosin VIIb knockdown tissues on the other hand appear indistinguishable from control.

**Figure 4:**
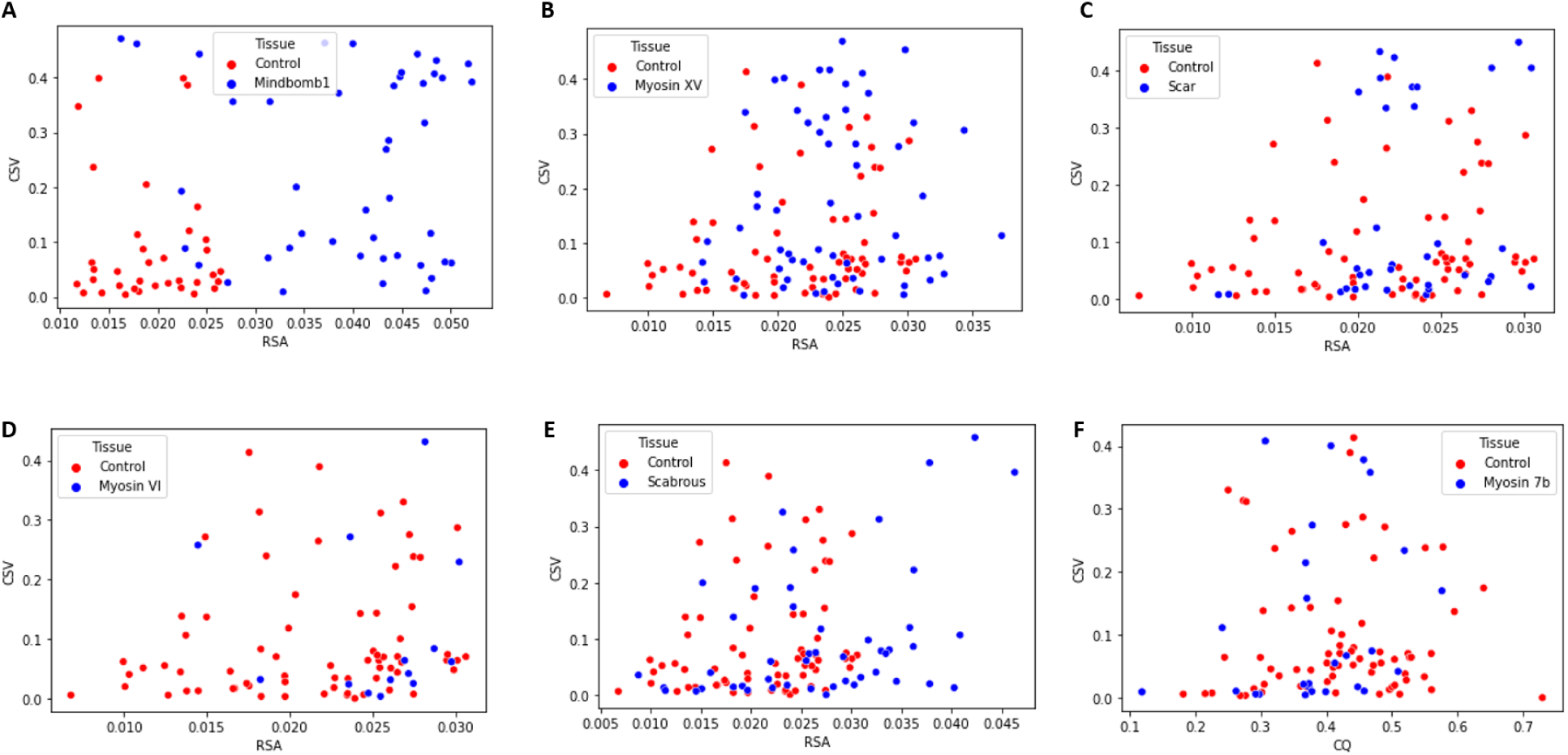
Scatter plot of significant features of tissue patterns. Distribution of a) Mindbomb1, b) Myosin XV, c) Scar, d) Myosin VI, e) Myosin VIIb and f) Scabrous pattern features. Control for Mindbomb1 is Dcr2 while control for other mutants is White.

In order to determine a range of wild type patterning, we sought an optimal decision boundary that can separate all controls (control-White and control-Dcr2) from mutant patterns that show the clearest separation from control (Mindbomb1, Myosin XV and SCAR). Support vector machines are well suited for such classification tasks. We used a grid search over the common kernel type (linear, poly, rbf) and related hyperparameters to ensure an optimal decision boundary. The models were trained using cross validation on 60% of observations (145 patterns). See Materials and Methods for details on the grid search and training methodology.

The optimal decision boundary, obtained from an rbf kernel with *γ* = 0.001, is shown in Fig. 5A along with a scatter plot of the training data. The axes are scaled with respect to the mean (*µ*) and standard deviation (*σ*) of control-White features given in Table. 1. Notice an almost linear separation between wild-type and mutant patterns, with the region of wild type patterning defined by the purple triangular region below the decision boundary. Within this region we observe that majority of wild type patterns have their density within the range of [*µ_RSA_* −2∗*σ_RSA_, µ_RSA_* +*σ_RSA_*], with a cluster size variance in the interval [*µ_CSV_* − *σ_CSV_, µ_CSV_* + <u>^1^</u> *σ_CSV_*]. The SVM decision boundary accurately predicts 70% of observations in the test data set (98 patterns). About 73% of wild type patterns are identified to lie within the normal region while 68% of mutant patterns lie outside this region.

**Figure 5:**
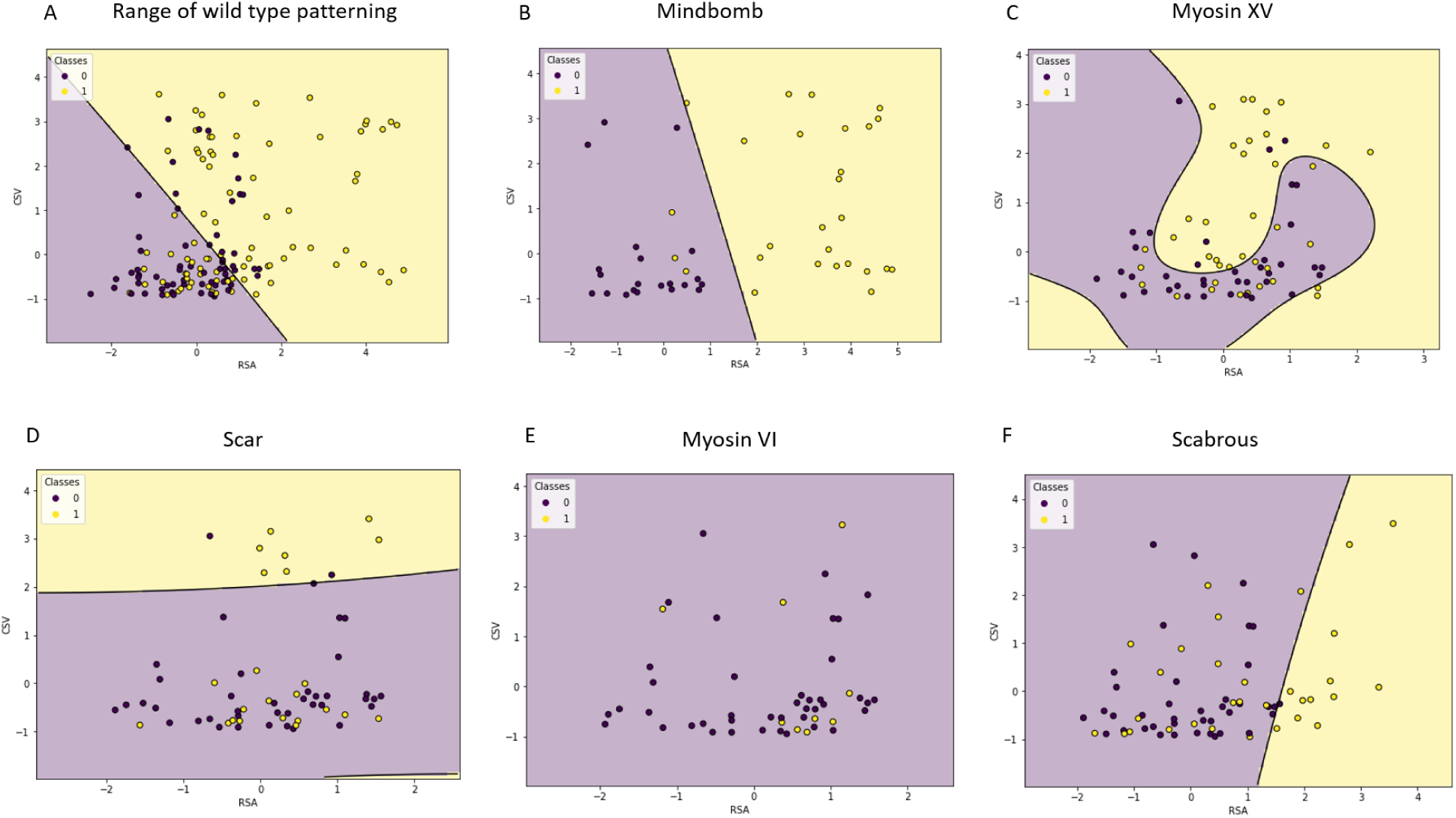
SVM decision curves for select mutant tissues based on RSA and CSV featuers. Scatter plot and decision curves showing a) range of wild type:0 (control White and control Dcr2) patterns compared with mutant:1 (Mindbomb, Myosin XV and SCAR mutant) patterns. Separation of b) Mindbomb:1 and Control Dcr2:0, c) Myosin XV:1 and Control White:0, d) Scar:1 and Control White:0, e) Myosin VI:1 and Control White:0, f) Scabrous:1 and Control White:0. All features are normalized using the mean and standard deviation of control-White presented in Table 1, prior to training.

### 3.4 Influence of individual genetic mutations on bristle organization

Finally, we wanted to quantify the proportion of each knockdown tissue that is considered to lie within the wild type region. This is a measure of how effective a particular knockdown condition is in disrupting the wild type pattern as well as how widespread its effect is among knockdown tissues. To achieve this, we constructed SVM decision boundaries for scatter plots of each knockdown condition and its corresponding control. The results, shown in Fig. 5B-F, reveal that Mindbomb1 knockdown has the strongest effect on the pattern, mostly increasing its density. The SVM decision boundary accurately distinguishes 91% of the test data with 82% of knockdown tissues considered different from Dcr2 control tissues (Fig. 5B). In Myosin XV knockdown conditions, a little over half of tissues (55%) are considered different from control (Fig. 5C). SCAR knockdown tissues show a weaker separation with only about 31% considered different from control tissues (Fig. 5D). Myosin VI knockdown patterns are completely indistinguishable from control (Fig. 5E), however Scabrous knockdown patterns show a better separation with 31% considered different from control (Fig. 5F).

## 4 Discussion

Our work examines five global features for quantifying the organization of spot patterns applied to analyzing sensory bristles on the dorsal thorax of *Drosophila melanogaster*. Among these, cluster number was found to be uninformative in distinguishing between wild-type and patterns from gene knockdown conditions. Our measure of global pattern density (RSA) confirms the known effect of Mindbomb1, Myosin XV, and Scabrous perturbation on the density of sensory bristles. Although SCAR is known to influence cytoneme formation, its density was not observed to be significantly different from control. Our analysis shows that Myosin VI regulates pattern density, with its knockdown condition showing significantly higher density of bristles. As mentioned earlier, Myosin VI motors are the only minus end directed actin motors with the potential to actively deliver Notch intracellular domain (NICD) activated along cytonemes to the cytoplasm. Genetic screens indicate Notch signaling is modified by decreased Myosin VI expression, and proteomics approaches have identified proteins in the Notch signaling pathway bind directly to the motor [36, 13, 37]. Since NICD translocation to the nucleus is a necessary step for Notch-mediated lateral inhibition, disruption of Myosin VI-mediated trafficking of NICD or other signaling components could increase the density of sensory bristles through reducing the efficacy of long-range signaling. No other gene considered significantly alters the density of sensory bristles.

In addition to increasing the bristle pattern density, our study reveals that Mindbomb1 knock-down patterns form clusters with more heterogeneous size than control and have a significantly smaller proportion of bristle precursor cells distributed further away from neighboring clusters. Mindbomb1 is a ubiquitously expressed E3 ubiquitin ligase that contributes to Epsin-mediated endocytosis of Delta ligand. Loss of Mindbomb1 activity leads to increased surface levels of Delta ligand, and therefore is linked to cis-inhibition of Notch receptor [38]. A model of macrochaete (large sensory bristles) selection hypothesizes that increased cis-inhibition leads to more dense bristle pattern [39]. It is unclear if microchaete are selected from proneural clusters in the same manner, or the extent to which cis-inhibition plays a role in this process. Other models that do not incorporate or have decreased long-range signaling also lead to dense patterns [6, 8]. Our results suggest that Mindbomb1 knockdown is also associated with more tissue-scale pattern variability from animal to animal. Notch signaling is used iteratively to arrange the five rows of bristles that extend along the anterior-posterior axis of each heminotum [4]. During these stages, Notch signaling is activated in large regions of nota tissue to define each bristle row. The role of Mindbomb1 during early pupal stages is not currently known, although its expression is required in the notum region of the developing wing disc during larval stages [29]. It is possible that the patterning variability we observed is due to an early notum requirement for Mindbomb1 in setting up tissue-scale organization, like bristle rows, in addition to within row patterning at later stages.

Myosin XV knockdown patterns also show more heterogeneity in the size of bristle precursor cell clusters. This result is consistent with the observation that Myosin XV knockdown leads to both increased pattern density as well as pattern disorganization [10]. Myosin XV is implicated in sub-cellular Notch and Delta localization, and disruption of ligand and receptor localization may lead to heterogeneity in long-range lateral inhibition. SCAR patterns are seen to form better clusters than control. However, one caveat is the lethality associated with RNAi-mediated knockdown of SCAR. Whereas Myosin XV is lethal at later pupal stages and Mindbomb1 may have overlapping function with other E3 ubiquitin ligases, SCAR mutants are lethal prior to pupal stages [40]. RNAi expressing animals that survive to pupariation likely have sufficient SCAR protein levels, leading to a mitigated patterning phenotype. Experiments previously investigating the role of SCAR in bristle patterning were performed using mutant tissue clones to avoid this issue [9, 5].

The process of bristle precursor fate selection is stochastic, leading to adult flies with similar, but not identical, bristle arrangements. This leads to a range of patterns that can be defined as wild-type, which could overlap to a varying extent with mutant or knockdown pattern phenotypes. Using the pattern density and cluster size variance, we were able to identify a region of wildtype patterning, bounded by an optimal svm decision boundary, which identifies 73% of wild-type patterns. It is possible that including more as yet unidentified parameters would increase the identification rate of wild-type patterns, however, intrinsic and environmental variability during development will likely still produce some patterns that will not be identified correctly. There are additional issues that can affect the rate of identification for gene knockdown conditions. Many factors contribute to the efficacy of an RNAi construct to knock down expression of its target. This may contribute to the penetrance of a phenotype, or the proportion of all RNAi-mediated knockdown tissues that exhibit a bristle patterning phenotype. RNAi efficacy can also contribute to the severity of the patterning phenotype.

Constructing decision boundaries for individual knockdown conditions, we found that Mind-bomb1 knockdown significantly disrupts the organization of 82% of patterns. This suggests a very strong penetrance for the patterning phenotype. This is not surprising since Mindbomb1 regulates ligand endocytosis, a necessary mechanism for Notch activation. The effects of other knockdown conditions are less dramatic, with Myosin XV knockdown having the second greatest effect (55% penetrance), followed by SCAR and Scabrous knockdown both at 31% penetrance. Surprisingly, all Myosin VI knockdown patterns were classified as control even even though our statistical analysis showed that the pattern density differed significantly. There is evidence that SCAR and Myosin XV influence cytoneme dynamics [10, 5], whereas Myosin VI could potentially affect transport of Notch signaling components along cytonemes. Scabrous, which may be distributed locally by cytonemes, is known to bind Notch receptor directly and promote signaling [41, 42, 43, 44]. Our results suggest that knockdown conditions that affect cytoneme dynamics has a stronger influence on pattern density and cluster heterogeneity compared to those with the potential to disrupt the dynamics of Notch trafficking along cytonemes.

This study limits the description of spot patterns to density and clustering features. Whereas these are reasonable for describing bristle precursor cell organization, it is by no means an exhaustive list. In particular, patterns that were found to be indistinguishable from control are only so based on the features we have considered and may differ for other features. Therefore, the search for distinguishing features of spot patterns need not stop here. This work provides a machine learning framework within which any new feature can be investigated.

## 5 Author Contributions

EA and GH conceived and oversaw this study. The manuscript was written by WA, GH, and EA. AN and GH performed the fly crosses and microscopy. SH processed microscopy images and extracted cell centroids for clustering. OW, WA, EA performed cluster analysis, statistical analysis and training of support vector machine.

## 6 Competing Interests

Authors declare no competing interests

## 7 Funding

OW and AN were supported by National Science Foundation’s Research Experience for Undergraduates (NSF REU) program, Award No. 2150280. GH is supported by NIH R35GM150782. EA, WA, SH are supported by the Joint NSF/NIGMS Mathematical Biology Program, R01GM152810.

## Notes

### Competing Interest Statement

The authors have declared no competing interest.

